# Critical role for P53 in regulating the cell cycle of ground state embryonic stem cells

**DOI:** 10.1101/674960

**Authors:** Menno ter Huurne, Tianran Peng, Guoqiang Yi, Guido van Mierlo, Hendrik Marks, Hendrik G. Stunnenberg

**Author notes:** Corresponding author: Hendrik G. Stunnenberg. These authors contributed equally (Menno ter Huurne, Tianran Peng).

## Abstract

Mouse Embryonic Stem Cells (ESCs) grown in serum-supplemented conditions are characterized by an extremely short G1-phase due to the lack of G1-phase control. Concordantly, the G1-phase-specific P53-P21 pathway is compromised in serum ESCs. Here we provide evidence that P53 is activated upon transition of serum ESCs to their pluripotent ground state using serum-free 2i conditions and modulates G1-phase progression. Our data shows that the elongated G1-phase characteristic of ground state ESCs is dependent on P53. RNA-seq and ChIP-seq analyses reveal that P53 directly regulates the expression of the Retinoblastoma (RB) protein and that the hypo-phosphorylated, active RB protein plays a key role in G1-phase control. Our findings suggest that the P53-P21 pathway is active in ground state 2i ESCs and that its role in the G1-checkpoint is abolished in serum ESCs. Taken together, the data reveals a mechanism by which inactivation of P53 can lead to loss of RB and uncontrolled cell proliferation.

## Introduction

Mouse Embryonic Stem Cells (ESCs) are pluripotent and self-renewing cells derived from the inner cell mass of the mouse blastocyst. ESCs can be indefinitely maintained *in vitro* in serum medium supplemented with the cytokine Leukemia Inhibitory Factor (LIF)^1^ hereafter called serum ESCs. In the last decade, new serum-independent culture conditions have been developed^2,3^. Subsequent studies in the epigenome and transcriptome have revealed that the different culture conditions give rise to different flavors of ESCs that are thought to reflect developmental states^4,5^. Mouse ESCs cultured in chemically defined 2i medium (N2B27 neural differentiation medium supplemented with PD0325901, CHIR99021 and LIF, hereafter called 2i ESCs)^3^ were shown to have an unrestricted developmental potential and are therefore hypothesized to represent the ground state of pluripotency^4,5^.

The cell cycle of ESCs cultured in the presence of serum and LIF is extremely short, mainly due to truncated Gap-(G-) phases. The short G1-phase was considered to be characteristic of pluripotent mouse ESCs^6^. We have previously shown that the short G1-phase is characteristic of serum ESCs and is the result of ERK signaling. The latter pathway is inhibited in ground state pluripotent ESCs cultured in 2i resulting in an elongated G1-phase^7^. Proteins that delay G1 progression, e.g. the CDK2-inhibitors P21 and P27, are not expressed in serum ESCs^5,7–9^ but can be detected in 2i cultured G1-phase ESCs and contribute to the elongation of G1-phase. The combined knock out of P21 and P27 causes a decrease in G1-phase cells in 2i ESCs^7^. P21 and P27 prevent CDK-mediated phosphorylation and inactivation of the pocket proteins and thereby activate the G1-checkpoint.

Bypass of the G1 checkpoint in serum ESCs has been attributed to high levels of the CDK2 activator CDC25A^10^ and to the lack of a P53-mediated DNA damage response^11–13^. The observation that P21, a prominent target of P53 in G1-arrest^14^ and a ‘readout’ of P53 activity, is highly expressed in 2i and absent in serum ESCs suggests that the role or activity of P53 may be different^7^. *In vivo* studies indicate that P53 is active in the Inner Cell Mass (ICM) during early embryonic development^15^ and by extrapolation in ground state pluripotent cells that are most reminiscent of ICM. These observations are in line with growing evidence that P53 plays an important role in embryonic development and differentiation. The exact role of P53 in ground state ESCs is however still unclear.

Therefore, we set out to decipher the distinct roles of P53 in ground state 2i and serum conditions. We generated a P53 knockout in an ESC cell line expressing the FUCCI reporters that allow the designation of cells throughout the different phases of the cell cycle and subsequent analysis of specific populations. Our data show that P53 plays a critical role in G1-phase progression in ground state 2i-as compared to serum ESCs. Moreover, genome-wide P53 binding and the transcriptome of P53^-/-^ 2i ESCs reveals that P53 directly regulates Rb1 expression in ground state ESCs which affects the G1-phase.

## Results

### P53 regulates G1-phase progression in 2i ESCs

As a guardian of the genome, P53 minimizes the acquisition of DNA damage and plays a key role in maintaining genomic integrity in cells. A major pathway employed by P53 to prevent DNA damage is by halting G1-phase progression and S-phase entry via promoting Cdkn1a (coding for the P21 protein) expression, which results in the inhibition of the CYCLIN/CDK complexes^16^. The elevated expression of P21 and elongated G1-phase in 2i ESCs^7^ led us to hypothesize that P53 is active in 2i ESCs, but not in serum ESCs, and contributes to cell cycle regulation in the pluripotent ground state. Although the P53 protein level is surprisingly similar in these two ESCs states (Figure 1A), the proteomic analysis of chromatin associated^17^ and quantification of P53 protein levels in several cellular fractions indicated that the level of chromatin-bound P53 is slightly higher in 2i ESCs when compared to serum ESCs (Figure 1B).

**Figure 1:**
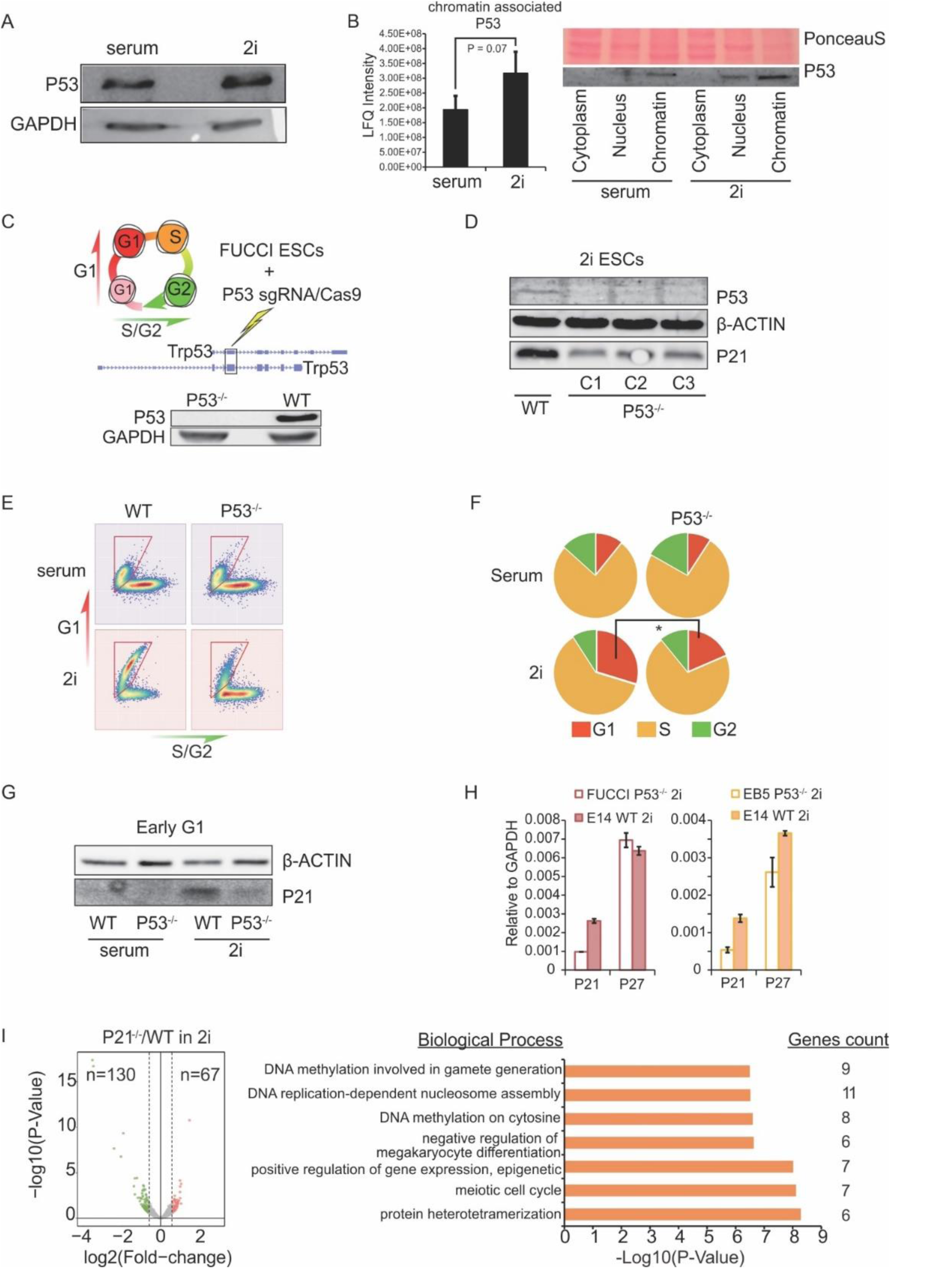
P53 is essential for the elongated G1-phase in 2i ESCs. **(A)** Western blot (WB) of P53 level in total cell lysate of wildtype serum and 2i ESCs. **(B)** Proteomics analysis and WB from different cellular fractions showing a higher level of chromatin-associated P53 protein in 2i ESCs as compared to serum ESCs. **(C)** Cells in G1-phase expressing the FUCCI reporter are Kusabira Orange-positive while cells in G2-phase are Azami Green-positive. Schematic representation of CRISPR/Cas9 mediated P53 knockout in FUCCI reporter ESCs targeting the common exon in Trp53 isoforms, resulting in the absence of P53 as shown on the Western Blot. **(D)** Three independent P53^-/-^ clones were obtained all showing a significant decrease in the expression of P21. **(E)** Analysis of FUCCI reporter expression in WT and P53^-/-^ cells. The longer G1-phase in 2i conditions (when compared to serum) is abbreviated in P53^-/-^ 2i ESCs, whereas in serum ESCs no differences between WT and P53^-/-^ ESCs are observed. Reporter expression is representative for three independent P53^-/-^ clones in at least two independent experiments. **(F)** Quantification of the different phases of the cell cycle in WT and P53^-/-^ EB5 cells using Bromo-deoxy-Uridine (BrdU) incorporation combined with Propidium Iodide (PI) staining. Significance was assessed using a two-tailed students T-test, *P-value < 0.05. **(G)** WB showing decreased expression of P21 during G1-phase in P53^-/-^ ESCs in 2i ESCs. **(H)** RT-qPCR reveals a reduction in P21 RNA levels in P53^-/-^ FUCCI ESCs and in an independent P53^-/-^ ESC line when compared to WT ESCs. No decrease in P27 mRNA was observed. RNA expression for all cell lines was measured in duplicate. **(I)** Volcano plot showing transcriptome changes in P21^-/-^ G1-phase ESCs compare to the WT G1-phase ESCs cultured in 2i conditions. Each dot represents one gene. Significantly changed genes were colored (down regulated genes in green and up regulated genes in red). GO clusters show the biological processes significantly enriched among the differential genes.

To determine the effect of P53 on the cell cycle of ESCs, we created three independent P53^-/-^ clones in R1 ESCs that express the FUCCI reporter constructs using the CRISPR/Cas9 gene editing system. The sgRNA were designed to cut the longest common exon of different P53 isoforms (Figure 1C). Upon deletion of P53, a clear reduction in P21 expression was observed in 2i cells (Figure 1D). As we reported previously, the proportion of the wild-type 2i ESCs in late G1-phase is much higher as compared to WT serum ESCs^7^. Therefore, we asked whether the G1-phase in P53^-/-^ ESCs is perturbed due to the decrease of P21. The FACS analysis of the P53^-/-^ cells showed a dramatic decrease in the number of 2i ESCs in late G1-phase (Figure 1E). Serum ESCs enter S-phase prematurely and therefore lack cells in late G1-phase. Accordingly, in P53^-/-^ serum ESCs virtually no effect on the cell cycle was observed. We next made use of an independent P53^-/-^ cell line (EB5, kindly provided by Hitoshi Niwa from Kumamoto University) to assess the distribution of cells over the different phases of the cell cycle using BrdU / PI staining. In 2i conditions, the number of cells in G1-phase was significantly lower in P53^-/-^ as compared to wild-type ESCs, further confirming our previous observations. Loss of P53 in serum ESCs had no measurable effect on the cell cycle (Figure 1F). Because P21 is primarily expressed during G1-phase in 2i ESCs^7^, its decreased expression in P53^-/-^ ESCs could be the result of the diminished number of cells in G1-phase. A western blot on G1-phase sorted cells shows that the expression of P21 is lowered specifically in 2i G1-phase cells (Figure 1G). Our previous study showed that deletion of P21 is not sufficient to significantly shorten the G1-phase, but requires the deletion of both P21 and P27^7^. However, the P27 expression level was either not affected (R1-FUCCI P53^-/-^) or slightly increased (EB5 P53^-/-^ cells) (Figure 1H). Furthermore, transcriptome analysis on WT and P21^-/-^ G1-phase ESCs displayed only mild changes in the gene expression with ∼130 were decreased and 67 were increased genes. The most significantly overrepresented biological processes were protein hetero-tetramerization, meiotic cell cycle and DNA replication-dependent nucleosome assembly^18,19^ (Figure 1I).

Taken together, we show that the P53 deficiency accelerates the G1-phase in 2i cells while no clear effect was observed in serum ESCs. Although P21 is downregulated upon deletion of P53, this alone is not sufficient to abbreviate the G1-phase, indicating that P53 may regulate the cell cycle independent of P21.

### Genes involved in cell cycle control are affected upon deletion of P53 in 2i ESCs

To determine the impact of P53 depletion on the transcriptome of serum- and 2i ESCs, we carried out RNA-seq on G1-phase sorted WT and P53^-/-^ ESCs in both culture conditions. The principal component analysis (PCA) plot shows that the variation in transcriptome between WT and P53^-/-^ is larger in 2i conditions as compared to serum conditions (Figure 2A; PC2). Differential expression analysis identified 1,430 significantly differentially expressed (DE) genes in 2i WT versus P53^-/-^, while only 321 DE genes were found in serum WT versus P53^-/-^ (Fold Change >1.5, adj. P-value <0.1) (Figure 2B). Over half (175 of 321) of DE genes in serum were also found to be differentially expressed in 2i ESCs (Figure 2C). In addition, the fraction of genes differentially expressed between WT and P53^-/-^ was higher in 2i as compared to serum (Figure 2D), suggesting a more extensive role of P53 in 2i conditions.

**Figure 2:**
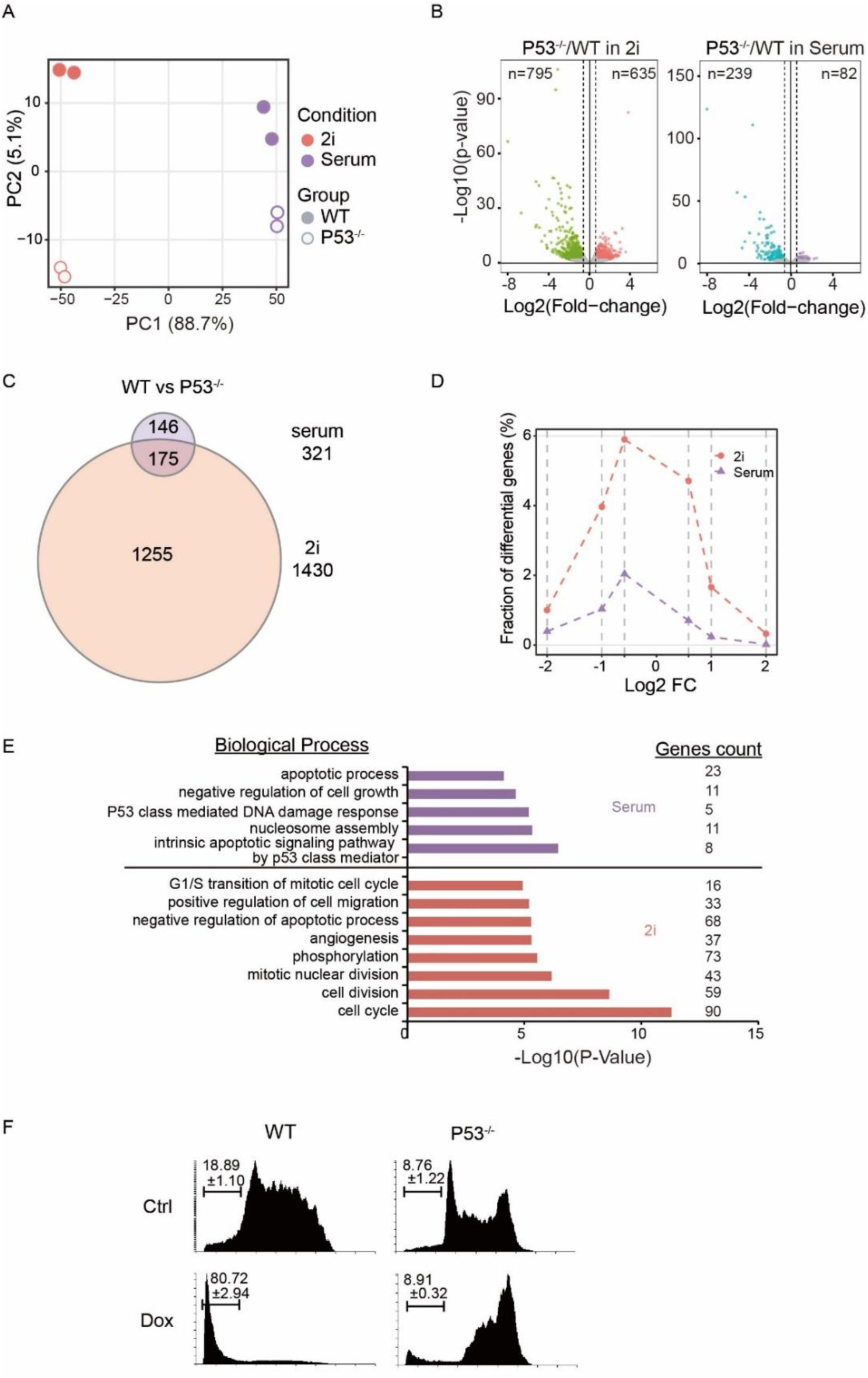
Genes differentially expressed in P53^-/-^ ESCs. **(A)** Principle Component Analysis on gene expression of WT and P53^-/-^ G1-phase ESCs cultured in serum and 2i conditions. **(B)** Volcano plot showing the transcriptome changes in 2i or serum P53^-/-^ G1-phase ESCs compared to WT. Each dot represents one gene. Significantly changed genes are colored (downregulated genes in green/blue and upregulated genes in red/purple). **(C)** Venn diagram showing the overlap of DE genes in serum and 2i. **(D)** Fraction distribution showing the fraction of genes differentially expressed between WT and P53^-/-^ ESCs cultured in either serum or 2i. **(E)** Gene Ontology analysis revealing the significant biological processes in serum or 2i ESCs. Number of genes present for each category are indicated at the right **(F)** Serum WT and P53^-/-^ ESCs treated with or without doxorubicin (1μM, 16 hours), stained by Propidium iodide and analyzed on FACS to assess apoptosis. Experiment performed in triplicate, numbers indicate mean percentage apoptotic ESCs (sub G1-phase) plus standard deviation. At least two independent experiments showed similar results.

To gain a deeper understanding of the biological processes involving P53 in serum as well as in 2i we performed GO analysis on DE genes (Figure 2E). In both serum and 2i conditions, genes downregulated in the P53^-/-^ cells are enriched for apoptosis-related processes. In line with these findings, the loss of P53 prevents apoptosis in both serum and 2i conditions upon doxorubicin treatment (1μM, 16 hours) as evident from the dramatic decrease in the number of cells in sub-G1-phase. In contrast to WT cells, the majority of P53^-/-^ cells stall in G2-phase after treated with doxorubicin (Figure 2F for serum conditions; 2i conditions are not shown) which is in line with recent findings showing that the loss of P53 does not affect doxorubicin-induced G2/M arrest but can abolish apoptosis of both primed and naive-state ESCs^20^. The analysis of the DE genes furthermore revealed that the set of genes differentially expressed 2i conditions is most significantly enriched for genes involved cell cycle processes, which is not observed for serum cultured ESCs. Thus, genes involved in the cell proliferation were highly affected in the G1 cells of 2i ESCs due to loss of P53.

### P53 activates Rb1 to elongate G1-phase in 2i ESCs

The differences in cell cycle and transcriptome between P53^-/-^ and WT ESCs indicate that P53 is essential for the elongated G1-phase in 2i conditions as compared to serum ESCs. Although there was a substantial decrease in the expression of P21 in the P53^-/-^ ESCs, the sole loss of P21 cannot explain the changes in the cell cycle^7^. Besides Cdkn1a (P21), numerous genes involved in cell cycle regulation are differentially expressed between P53^-/-^ and WT in 2i-cultured cells (Figure 3A). To identify direct targets of P53 connected to the cell cycle control, we next performed P53 ChIP-seq in serum and 2i ESCs. Supporting our prior observations that P53 has a more prominent role in 2i conditions as compared to serum conditions, the number of P53 binding sites is significantly higher in 2i (3,595 versus 1,347 in serum). To identify the genes that are likely under direct regulation of P53, we annotated the P53 ChIP-Seq binding sites using HOMER^21^. The result shows that 386 genes with P53 binding in their promoters or enhancers are also differentially expressed in 2i P53^-/-^ ESCs (Figure 3B). Out of these 386 genes 129 have a higher P53 ChIP-seq signal in serum (cluster 1) whereas 257 have a higher ChIP-seq signal in 2i (cluster 2).

**Figure 3:**
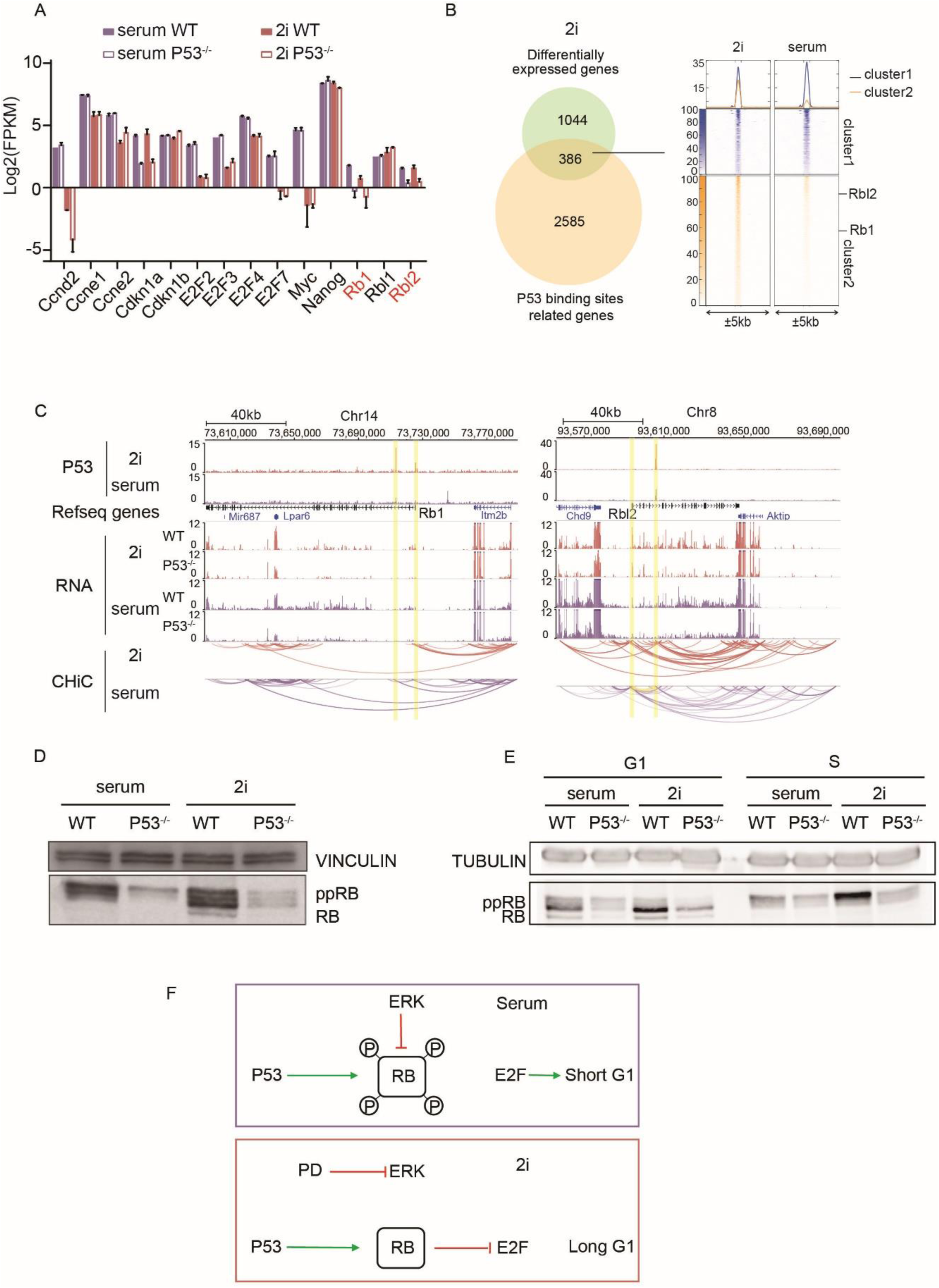
P53 activates Rb1 to elongate G1-phase in 2i ESCs. **(A)** Bar graph showing differential expression of cell cycle and other regulators in serum and 2i cultured WT and P53^-/-^ ESCs during G1-phase. The expression of Rb1, Rbl2 and Cdkn1a (P21) decreased in P53^-/-^ both in serum and 2i ESCs. **(B)** 2585 genes are associated with P53 binding sites, 386 of these genes are expressed differentially in 2i P53^-/-^ as compared to 2i WT cells **(C)** ChIP-seq and Capture HiC indicate that P53 binds regions that locate or interact with the Transcriptional Start Site (TSS) of Rb1 and Rbl2. **(D)** Western blot (WB) showing that the protein level of hyper-phosphorylated RB and hypo-phosphorylated RB is reduced in P53^-/-^ ESCs in serum and 2i, respectively. **(E)** WB showing the protein level of RB throughout the cell cycle in P53^-/-^ and WT ESCs in serum and 2i conditions. At least two independent experiments showed similar results. **(F)** Proposed model that displays the role of P53 and Rb1 in ESC cell cycle control.

Subsequent GO-term analysis revealed that this set of 386 genes was significantly enriched for genes that regulate cell cycle arrest (10 genes with a Benjamini corrected P-value of 9.6e-3). Interestingly, Rb1 is among those genes. We found that in P53^-/-^ ESCs the Rb1 and Rbl2 transcripts that encode RB and P130, respectively, are decreased as compared to WT ESCs (Figure 3A). The pocket proteins RB and P130 are well known to be involved in the control G1-phase progression in 2i ESCs by inhibiting the activity of the E2F transcription factors^7^ and are themselves downstream targets of the CDK/CYCLIN pathway. P53 ChIP-seq in 2i ESCs showed that P53 binds to the Transcriptional Start Site (TSS) and gene body of Rb1 and to the gene body of Rbl2. Integrating analysis of RNA-seq, ChIP-seq and Capture HiC data^22^ revealed that the P53 binding sites interact with the TSS of Rb1 and Rbl2 suggestive of a direct transcriptional regulation (Figure 3C).

Although the expression of these pocket proteins was drastically reduced both in serum as well as in 2i conditions after P53 knockout, inactive hyper-phosphorylated RB was the predominant form in serum ESCs, whereas hyper- as well as active hypo-phosphorylated RB could be detected in 2i ESCs (WT was shown previously^7^) (Figure 3D). Since the cell cycle distributions of asynchronously growing WT and P53^-/-^ 2i ESCs are different, we next determined RB protein levels throughout the cell cycle in WT and P53^-/-^ serum and 2i ESCs. The result showed a clear reduction of RB expression in P53^-/-^ ESCs and furthermore showed that the fraction of hypo-phosphorylated RB was higher in 2i than in serum G1 cells (Figure 3E).

Taken together, our data strongly suggests that P53 directly activates the transcription of the pocket proteins RB and P130 thereby elongating G1-phase in 2i ESCs. Due to increased ERK-/CDK/CYCLIN-signaling RB is constitutively hyper-phosphorylated in serum ESCs and the deletion of P53 has minor effect in these cells (Figure 3F).

## Discussion

Rapid proliferation is a hallmark of pluripotent stem cells and has intrinsically been associated with their unique cell cycle^23^. A truncated G1-phase is fundamental to the cell cycle of ESCs and is reflected by their smaller size when compared to somatic cells^24^. The shortened G1-phase is accompanied by the impairment of pathways that control genomic integrity during G1-phase in serum ESCs^25^. However, the long-standing dogma that ESCs have a shorter G1-phase than somatic cells was mainly based on studies performed in serum-cultured ESCs^9,26^. Recently, we and others have shown that 2i ESCs have a much longer G1-phase as compared to serum ESCs^7,27^. How these differences in cell cycle between serum and 2i ESCs affect the biological processes that take place during G1-phase has remained unclear.

P53 is well known for its pivotal role in induction of G1-arrest to protect genomic integrity. Early studies in serum ESCs have found that although P53 is highly expressed, it cannot act as a regulator of G1-phase progression due to functional uncoupling of the P53/P21 axis^28^. The elevated expression of P21 and the identification of the elongated G1-phase led us to hypothesize that P53/P21 pathway maybe activated and extend the G1-phase upon transition of serum ESCs to 2i conditions. The results presented here indicate that P21 expression is indeed regulated by P53. Unexpectedly, however, the elongated G1-phase in 2i ESCs depends on a novel unexplored function of P53 in the cell cycle of ESCs. In ESCs, P53 regulates not only the expression of P21 but also that of the downstream pocket proteins RB and P130. In serum, but not in 2i ESCs, the pocket proteins are inactivated due to abundant ERK signaling^7^, explaining why this function of P53 has not been observed in serum ESCs. By regulating the expression of the pocket proteins, P53 is crucial for the elongated G1-phase in 2i conditions. The loss of P53 does, however, not fully shorten the G1-phase in 2i ESCs to the level of that in serum ESCs, which suggests other mechanisms are involved as well in regulation of G1-phase progression in 2i ESCs. The lowered ERK signaling and reinstatement of the P53-mediated G1-checkpoint in 2i ESCs suggests that these cells are better able to cope with DNA damaging events, which however remains to be shown.

P53 is highly expressed in the early embryo, but its functional role is still elusive. Our findings suggest that in the early embryo where ERK signaling is absent, P53 plays a critical role during G1-phase to restrict rapid cell proliferation by modulating the expression of the pocket proteins. Interestingly, the cellular senescent state that resembles diapause *in vivo* is depending on the presence of this family of proteins^7,29^. Our observations therefore suggest that P53 plays a role in diapause. Possibly, P53 is highly expressed in early rodent embryos in order to induce diapause in response to stressful conditions.

Besides the differences in cell cycle control between WT and P53^-/-^ 2i ESCs, the RNA-seq data uncovered a large number of developmental genes (amongst others involved in angiogenesis and the development of the nervous system) that are negatively affected by loss of P53 in 2i conditions. These findings are in line with previous reports that suggest an important role for P53 in differentiation, and imply that P53 is functionally more dynamic in 2i. How P53 regulates the expression developmental genes in 2i remains to be determined, possibly the differential regulation of the pocket proteins plays a role, considering their role in development and differentiation^30,31^.

Altogether we show that in ESCs the function of P53 differs depending on the cellular state. In ground state 2i ESCs P53 is involved in controlling the cell cycle via directly regulating the expression of the pocket proteins.

## Acknowledgment

We thank Onkar Kiran Joshi for offering the Capture HiC data and Haoyu Wu for the assistance of ChIP-seq. We thank Masaki Shigeta and Hitoshi Niwa for kindly providing the EB5 P53^-/-^ ESCs^32^. This work was supported by ERC grant ERC-2013-AdG No. 339431 – SysStemCell (to H.G.S.).

## Conflict of interest

The authors declare no conflict of interest

## Author Contributions

M.t.H, T.P and H.S.G conceived the paper and wrote the manuscript. M.t.H and T.P performed experiments. M.t.H, T.P and G.Y analyzed the data. G.v.M generated and analyzed the proteomics data. H.M contributed to the study design and helped drafting the manuscript. Funding was obtained by H.G.S.

## Materials and Methods

### Cell culture

Mouse ESCs were cultured on feeder-free 0.1% gelatin coated plates in a 37° humidified incubator with 5% CO_2_. Serum medium consists of DMEM with GlutaMax (Gibco #31966-021) containing 15% fetal bovine serum (Hyclone #SV30160.03), LIF (1000U/mL, Millipore #ESG1107), sodium pyruvate (1x, Gibco#11360-070) and β-mercaptoethanol (50 μM, Millipore#8.05740.0250). 2i medium consists of NDiff 227 (Takara #Y40002) supplemented with MEK inhibitor PD0325901 (1μM), GSK3 inhibitor CHIR99021 (3 μM) and LIF (1000U/mL). Cells were refreshed every day and passaged every 3 days.

### Immuno blotting

Cells were trypsinized and washed with PBS. Cell pellet were lysed in RIPA buffer (150mM NaCl, 1% NP-40, 0.5% NaDOC, 0.1% SDS, 50mM Tris-HCl pH=8) with fresh EDTA-free protease inhibitor cocktail (Roche #4693132001). Protein concentration was measured using Bio-rad protein assay (#500-0006). Cell extracts were loaded equally and separated by 7%∼12% SDS-PAGE, electrotransferred to nitrocellulose membranes and incubated in blocking buffer (5% nonfat milk in TBST) for 1 hour at room temperature. Membranes were incubated with primary antibody (1:1000 diluted in blocking buffer) over night at 4°C then washed 5 times in TBST for 5 minutes at room temperature and incubate with second antibody (1:2000 diluted in blocking buffer) at room temperature for 1 hour. After five washes with at room temperature for 5 minutes ECL substrate (ThermoFisher #32106) were added and images were acquired. The primary antibodies used in this study are P53 (Oncogene #OP03), P21 (SantaCruz #sc-6546), RB (BD Bioscience #554136), GAPDH 6C5 (Abcam #8245), TUBULIN (Santacruz, #sc5286), VINCULIN (Santacruz,#sc5573) The secondary antibodies used are Swine anti-Rabbit HRP (Dako #P0217), Rabbit anti-Rat HRP (Dako #P0450) and Rabbit anti-Mouse HRP(Dako #P0161).

### Genome editing using CRISPR-Cas9

We made use of the CRISPR-Cas9 gene editing technology to knock out Trp53 (P53). An online tool (crispr.mit.edu and Benchling) was used to design the gRNAs that were cloned into the plasmid Cas9(BB)-2A-GFP (Addgene plasmid 48138) as described previously^7^. FUCCI serum ESCs were transfected using lipofectamine-3000 (Thermofisher # L3000015). After 48 hours, cells expressing GFP over background (from the Azami Green reporter) were sorted with a BD FACS Aria. Cells were split at clonal density and after approximately 7 days colonies were picked for expansion. Genomic DNA from individual clones was extracted using the Wizard Genomic DNA extraction kit. The targeted region was PCR amplified and Sanger Sequenced. gRNA oligonucleotides were as follows:

Trp53_gRNA_a_Fwd CACCGGAGCTCCTGACACTCGG

Trp53_gRNA_a_Rev AAACCCGAGTGTCAGGAGCTCC

### Flow cytometry

The BD FACS Aria cell sorter was used to analyze and sort FUCCI ESCs. For FUCCI reporter quantification CSV files were analyzed using the Flowcore Package. For cell cycle analysis cells were pulsed with 20 µM BrdU for 30 minutes, harvested by trypsinisation and fixed overnight in 70% ethanol at 4°C. After denaturation of the DNA using 2N HCl + 0.5% Triton-X100 for 30 minutes at room temperature, neutralization with 0.1M Na2B4O7 (pH 8.5), samples were incubated with the anti-BrdU antibody over night at 4°C. Next day, the samples were stained with Propidium Iodide staining solution (10 ug/ml PI [Sigma, P4170] and 0.2mg/mL RNAse A in PBS) over night at 4°C. Samples of at least 10000 cells were acquired using a FACScalibur or FACSverse flow cytometre (Becton Dickinson). Subsequent analysis was done with Flowing Software.

### qRT-PCR and RNA-seq

Total RNA were extracted using the RNeasy Mini Kit (Qiagen #74106) following the manufacturer’s protocol. SuperScript™ III Reverse Transcriptase (ThermoFisher #18080093) and random primers (p(dN)_6_, Roche #11034731001) or Oligo(dT)_12-18_ primer were used for reverse transcription. Real-time qPCR was performed using the iQ™ SYBR® Green Supermix (Bio-Rad #1725006). An endogenous control (Gapdh, Forward: TTCACCACCATGGAGAAGGC, Reverse: CCCTTTTGGCTCCACCCT) was used to normalize the expression. Biological replicates were performed for all RT qPCR reactions. P21 and P27 primers have been described before^33^.

5 µg of extracted RNA was depleted from ribosomal RNA using Ribo-Zero Gold Kit (Epicentre Madison). After fragmentation of the rRNA-depleted RNA, 500ng was reverse-transcribed using SuperScript™ III Reverse Transcriptase and random primers (dN)_6_ following the manufacturer’s instructions. Next, libraries were prepared using the KAPA Stranded RNA-Seq Library Preparation Kit (KAPA #8400) following the manufacturer’s instructions.

### ChIP-seq

ESCs were fixed using 1% formaldehyde (Millipore #344198) for 10 min at room temperature then quenched by adding 1.25M glycine to a final concentration of 0.125M. Cell pellets were snap frozen and stored in −80°. Every 10 million cells were lysed by 300μL 1% SDS include freshly made Protease Inhibitor (PI) cocktail (Roche #4693132001), sonicated with Diagenode Bioruptor Pico and diluted with 2.7mL PBA (1x PBS + 0.5% BSA) with fresh added PI cocktail. Dynabeads protein A+G (Invitrogen #10008D, 10009D) were washed and pre-blocked with cold PBA for 30 minutes. Every 30μg diluted chromatin (∼250μL) were incubated with 15μg P53 antibody (Novocastra #CM5P) and 60μL pre-blocked beads and rotated at 4° overnight. After incubation, beads were washed with High-NaCl/Low-NaCl/LiCl/TE /TE washing buffer for 10 minutes and transferred to new eppendorf tube. Beads were eluted with 200µL elution buffer (1%SDS, 0.2M NaCl, 0.1μg/μL Proteinase K) at 65°C thermo shaker 1000rpm for 20 minutes. Supernatants were purified with MinElute PCR Purification Kit (QIAGEN #28006). 1-5ng of DNA was used for library construction with KAPA Hyper Prep Kit (KAPA #KK8504).

### ChIP-seq analyses

For each sample, all 42bp reads were mapped onto the mouse genome (mm9) using Burrows-Wheeler Aligner (BWA) aligner^34^ with default parameters. The mapped reads were regarded as input for Picard Mark Duplicates (http://broadinstitute.github.io/picard/) to remove potential PCR duplicates. MACS2^35^ was used to call P53 peaks with a narrow q-value cut-off of 0.01. Read density profiles are displayed as fold enrichment track generated by normalizing ChIP data over input DNA pileup signal files using MACS2. These profiles were further visualized by deepTools^36^.

### RNA-seq analyses

RNA sequencing reads were aligned to the mouse reference genome mm9 using STAR tool^37^ which could enumerate gene-level read counts at the same time. The differentially expressed genes were identified with the DESeq2 package^38^ by comparing knockout with wild-type groups. Only those genes greater than 1.5-fold changed at Benjamini-Hochberg-corrected P-value < 0.1 were considered significantly deregulated. The transcriptional levels of genes were estimated as Fragments Per Kilobase per Million aligned reads value (FPKM) values using by Cufflinks^39^.

### Dimensionality reduction and functional annotation

To explore potential variances between different groups, we performed principal component analysis. Top 3,000 variable genes were first selected based on interquartile range (IQR) of normalized gene expression levels, and further used to reduce dimensionality of the dataset by pca function in R. We used DAVID tool^19^ to assess enriched gene ontology terms and pathways in order to gain insight into the biological functions for deregulated genes. Only terms with Benjamini-adjusted P-value < 0.05 were considered significantly overrepresented.

